# High resolution and high throughput bacteria separation from blood using elasto-inertial microfluidics

**DOI:** 10.1101/2020.10.19.344994

**Authors:** Sharath Narayana Iyengar, Tharagan Kumar, Gustaf Mårtensson, Aman Russom

## Abstract

Improved sample preparation has the potential to address a huge unmet need for fast turnaround sepsis tests that enable early administration of appropriate antimicrobial therapy. In recent years, inertial and elasto-inertial microfluidics-based sample preparation has gained substantial interest for bioparticle separation applications. However, for applications in blood stream infections the throughput and bacteria separation efficiency has thus far been limited. In this work, for the first time we report elasto-inertial microfluidics-based bacteria isolation from blood at throughputs and efficiencies unparalleled with current microfluidics-based state of the art. In the method, bacteria-spiked blood sample is prepositioned close to the outer wall of a spiral microchannel using a viscoelastic sheath buffer. The blood cells will remain fully focused throughout the length of the channel while bacteria migrate to the inner wall for effective separation. Initially, particles of different sizes were used to investigate particle focusing and the separation performance of the spiral device. A separation efficiency of 96% for the 1 µm particles was achieved, while 100% of 3 µm particles were recovered at the desired outlet at a high throughput of 1 mL/min. Following, processing blood samples revealed a minimum of 1:2 dilution was necessary to keep the blood cells fully focus at the outer wall. In experiments involving bacteria spiked in diluted blood, viable *E.coli* were continuously separated at a total flow rate of 1 mL/min, with an efficiency between 82 to 90% depending on the blood dilution. Using a single spiral, it takes 40 minutes to process 1 mL of blood at a separation efficiency of 82% and 3 hours at 90% efficiency. To the best of our knowledge, this is the highest blood sample throughput per single microfluidic chip reported for the corresponding separation efficiency. As such, the label-free, passive and high throughput bacteria isolation method has a great potential for speeding up downstream phenotypic and molecular analysis of bacteria.

## Introduction

Sepsis is an acute inflammatory response to pathogens by an immune-compromised host body. Sepsis may be defined as the body’s systemic inflammatory response (SIRS) to an infection caused by pathogens. If left untreated, it may lead to shock, multi-organ failure, and death – especially if not recognized early and treated promptly. Sepsis is the final common pathway to death from most infectious diseases worldwide, including viral infections such as SARS-CoV-2 / COVID-19^1^. The definition of sepsis (Sepsis 3) was updated in 2016 and is currently defined as a life threatening condition caused by a dysregulated host response due to microbial infection leading to host organ dysfunction^2^. The term ‘septic shock’ was also defined in the same study as a subset of sepsis, where a substantial increase in mortality occurs due to profound cellular, metabolic, and circulatory abnormalities. The development of sepsis starts from a site of local infection where bacteria enter into the blood stream. The most common bacteria causing sepsis include *E.coli* and *Staphylococcus* among others such as *Pseudomonas, Klebsiella, Candida, Acinetobacter* etc ^3,4^. The detailed etiology of organisms causing sepsis highlighting the frequency of infection and their corresponding mortality rate has been studied and reported extensively ^3, 5^. According to 2018 statistics from the World Health Organization (WHO) ^6, 7^, about 30 million people are affected by sepsis worldwide, among which approximately 3 million are new born and 1.2 million are children. Of those affected, 6 million people die worldwide. More than 500 thousand new born babies die each year, and one in ten maternal death occur. A recent study reported that sepsis-related deaths globally are double to what was previously estimated ^8, 9^. Sepsis also remains the most common cause of in-hospital deaths in the United States, costing the country $24 billion a year in 2013^10^. In a majority of the cases, the causative pathogen is bacteria^3^. Rapid diagnostics and therapy administration play a vital role in sepsis-related mortality^11^. Sepsis can lead to septic shock, multiple organ failure, and ultimately death, with an associated rise in mortality of 7% for every hour delay in the administration of appropriate antibiotics^12^. Currently, the “gold standard” for diagnosis of blood stream infections (BSI) is by blood cultures. It can take 24-72h before a complete answer can be reached, including the antibiotic resistance profile ^13–15^. In addition, detection of fastidious bacteria and fungi, which often require longer time to grow, is challenging ^16^. For faster turnaround time, nucleic acid-based techniques (NAT) are promising, e.g. IRIDICA, SeptiFast, SeptiTest or U-dHRM ^17^. These kits combine lysis buffers for fast blood sample pre-treatment and DNA extraction, with PCR analysis enabling the detection of bacterial DNA, but with low sensitivity and specificity. This is mainly due to background from the human DNA ^17^. Although tremendous improvements have been made over the past few years, including the introduction of next generation sequencing technologies, sample preparation remains the bottleneck for further expansion of molecular diagnostics into the clinical settings.

Among emerging technologies, microfluidics is very promising tool and has the potential to enrich the pathogens from blood sample in a seemingly integrated fashion. The ability to precisely manipulate blood cells has attracted considerable research in the field and several attempts have been made to address sample preparation for sepsis diagnostics. Common methods either use the surface property to isolate bacteria using affinity separation ^18–21^, or other properties, such as their shape, size, deformability, density, electric or magnetic susceptibility, and hydrodynamic properties ^22–38^. In general, there is often a trade-off between efficiency and throughput. To this end, inertial microfluidics is attractive since the passive, size-based, technology exploits inherent hydrodynamic forces that scale with increased flow rate and typically operate at extremely high volumetric flow rates (∼mL/min). However, while the technology has been utilized extensively to precisely focus and separate mammalian cells including circulating tumor cells^39^, separation of bacteria has been challenging. Initial work in inertial microfluidics were mostly focused on separation of bacteria from diluted red blood cells ^28^. Using a spiral device, Hou et al. recently used a sheath flow at the inlet to push the blood cells toward one side of the inlet wall and performed size-based differential migration of blood cells to separate the bacteria ^37^. Here, the bacteria are smaller and lag behind the cells and could be extracted at an efficiency of 70%.

The main challenge in inertial microfluidics-based bacteria separation is due to the narrow size difference between bacteria (~1-2 µm) and blood cells (~ 3–15 µm). One way of improving the size resolution is to use the rheological properties of viscoelastic fluid itself to manipulate the cells. In elasto-inertial microfluidics, the combination of fluid elasticity and fluid inertia enable size-based focusing of particles ^40–42^. Using a straight rectangular channel, we recently reported the separation of bacteria from whole blood by selectively migrating blood cells away from the walls towards the center line of the channel while bacteria remain in the streamline and could be separated ^30^. While the method offers capabilities for precise cell manipulation, the relatively low separation efficiency and limited throughput is hindering practical implementation. The relatively low volumetric flow rate is an inherent limitation of elasto-inertial microfluidics since the synergetic effect of the elastic forces and inertial forces are ideal at moderate flow rates. In this work, using a spiral microchannel, we utilize viscoelastic flows at extremely high volumetric flow rate (∼mL/min).

Using PEO (Polyethylene Oxide) as an elasticity enhancer, we report on blood cell focusing and bacteria separation in spiral channels at extremely high volumetric flow rates (~ mL/min range). Using a two inlet and two outlet spiral device, diluted blood sample is pinched towards the outer wall by the PEO buffer. As the hydrodynamic forces develop, the blood cells start to be fully focused at the outer wall and the focusing position is maintained throughout the channel length, while smaller bacteria follow the Dean vortices and are effectively separated and collected at the inner outlet. First, the focusing and separation phenomena of the spiral device were studied using microparticles. Using 1μm particles as a model for bacteria, it was possible to separate the particles from 3 μm particles at a an extremely high flow rate (1 mL/min). Thereafter, the optimized spiral geometry was utilized to separate bacteria from diluted blood. First, blood cells spiked with 1μm particles were investigated to identify the optimal dilution conditions where the blood cells remain close to the outer wall while the 1μm particles readily migrate towards the inner wall for separation. Finally, using *E.coli* bacteria spiked into diluted blood sample as a model for sepsis, bacteria separation is demonstrated at an efficiency of 82 to 90% depending on the blood dilution (1:2 to 1:10 dilution).

## Theoritical background

A Poiseuille flow of an incompressible Newtonian fluid through a straight channel will exhibit a parabolic velocity flow profile with a maximum velocity at the center and zero velocity at the walls of the channel, i.e. a no-slip boundary condition. The flow of a fluid in the channel is characterized using a dimensionless number Reynolds number (Re), defined as 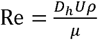,where 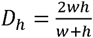 is the hydraulic diameter with ‘*w*’ and ‘*h*’ being the width and height of the channel respectively, ‘*U*’ is the bulk velocity of the fluid, ‘*ρ*’ is the fluid density and ‘*µ*’ is the dynamic viscosity. The equilibrium position of the particle suspended in the fluid is mainly due to the combined effect of shear induced lift force (F_LS_) and wall induced lift force (F_W_). The parabolic flow profile of the fluid induces F_LS_, pushing the particle away from the center of the channel while F_W_ pushes the particle away from the wall. The combination of these two forces, F_L_, defined as 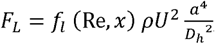, will result in an equilibrium focusing position of the particle. Here, *f*_*l*_(*R*_*e*_,*x*) is the lift coefficient, which is a function of Re, ‘*x*’ is the position of the particle and ‘*a*’ is the size of the particle ^22^. The number of equilibrium positions of the particles depend upon the geometry of the channel. Segre & Silberberg in 1961, showed that, particles attain their equilibrium positions along the annulus of a circular channel at 0.6 radius ^43^. In 2007, Di Carlo et al. observed that in a square channel, particles occupy equilibrium positions at the faces of each wall of the channel ^22^ while Bhagat et al. demonstrated equilibrium positions along the longer sides of a rectangular channel ^44^. In case of a fluid flowing through a curved channel, the curvature of the channel creates a radial pressure gradient creating secondary fluid flows leading to the formation of two counter-rotating Dean vortices, normal to the bulk flow. The drag force, F_D_, caused due to Dean vortices is defined as 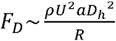^22^, where ‘*R*’ is the radius of curvature of the channel. The strength of the Dean vortices is determined by a dimensionless Dean number defined as 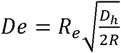 ^45^. In contrast to a Newtonian fluid, when a viscoelastic (non-Newtonian) fluid flows through a channel, an additional elastic force, F_E_, arises due to the first normal stress difference of the fluid which pushes the particle toward the center of the channel. This force is defined as where ‘*N*_*1*_’ is the first normal stress difference ^41, 46, 47^.The elastic property of the fluid is defined by the dimensionless Weissenberg number, —, where ‘*Q*’ is the volumetric flow rate of the fluid and ‘ is the relaxation time of the fluid ^48^.

In this paper, the complex interaction and balance between the above forces are utilized to demonstrate focusing and differential migration of bacteria from fully focused blood cells to achieve size-based separation at high flow rates.

## Results and discussion

### High throughput particle focusing

In viscoelastic flows through curving microchannels, the following dominant forces affect particle focusing: lift force (F_L_), elastic force (F_E_), and a curvature-induced Dean’s drag force (F_D_). The combined interaction results in particles migration and focusing. While previous work from our group and others have mainly focused on particle migration towards an equilibrium position using either inertial and elasto-inertial microfluidics, in this work we show that it is possible to pre-position particles at the inlet and keep the particles fully focused throughout the channel length strictly based on particle size. To this end, for the first time, we show that particles above a certain size cut-off, prepositioned at the lateral equilibrium position will remain fully focused throughout the channel, independent of length, in flow through spiral microchannels. Smaller particles are affected differently by the dominant forces and entrapped into the Dean vortices and continuously migrate away from the equilibrium position. By optimizing the spiral geometry and length, we show it is possible to fully migrate the smaller particles close to the inner wall for efficient separation By. Experimentally, by simply introducing a viscoelastic sheath buffer (PEO) into a two-inlet spiral microchannel, the blood will be pinched toward the outer wall. The pre-positioned blood cells will remain fully focused while smaller bacteria readily migrate
 towards the inner wall and can be extracted. Fig.1 shows a schematic illustration of how the method works. A sheath fluid pinches the sample containing large and small particles to occupy a narrow stream at the outer wall of the inlet. As the hydrodynamic forces are developed, the larger particles remain focused due to the balance between the three main forces while the smaller particles are trapped in the Dean vortices and differentially migrate away from the outer wall. Finally, the smaller particles reach the inner wall and can then be separated. Note, the total flow rate (sample + sheath) is extremely high, in the range of mL/min. Furthermore, the focusing phenomena differ from inertial microfluidics, where the focusing point is toward the inner wall laterally and at two vertically focusing points as a result of balance between FL and FD. We evaluated several parameters that influence the particle focusing behavior, preposition of the particle at the inlet, flow rate and particle size. To this end, we first used microparticles to examine the focusing phenomena before evaluating the spiral for bacteria separation.

**Fig. 1.**
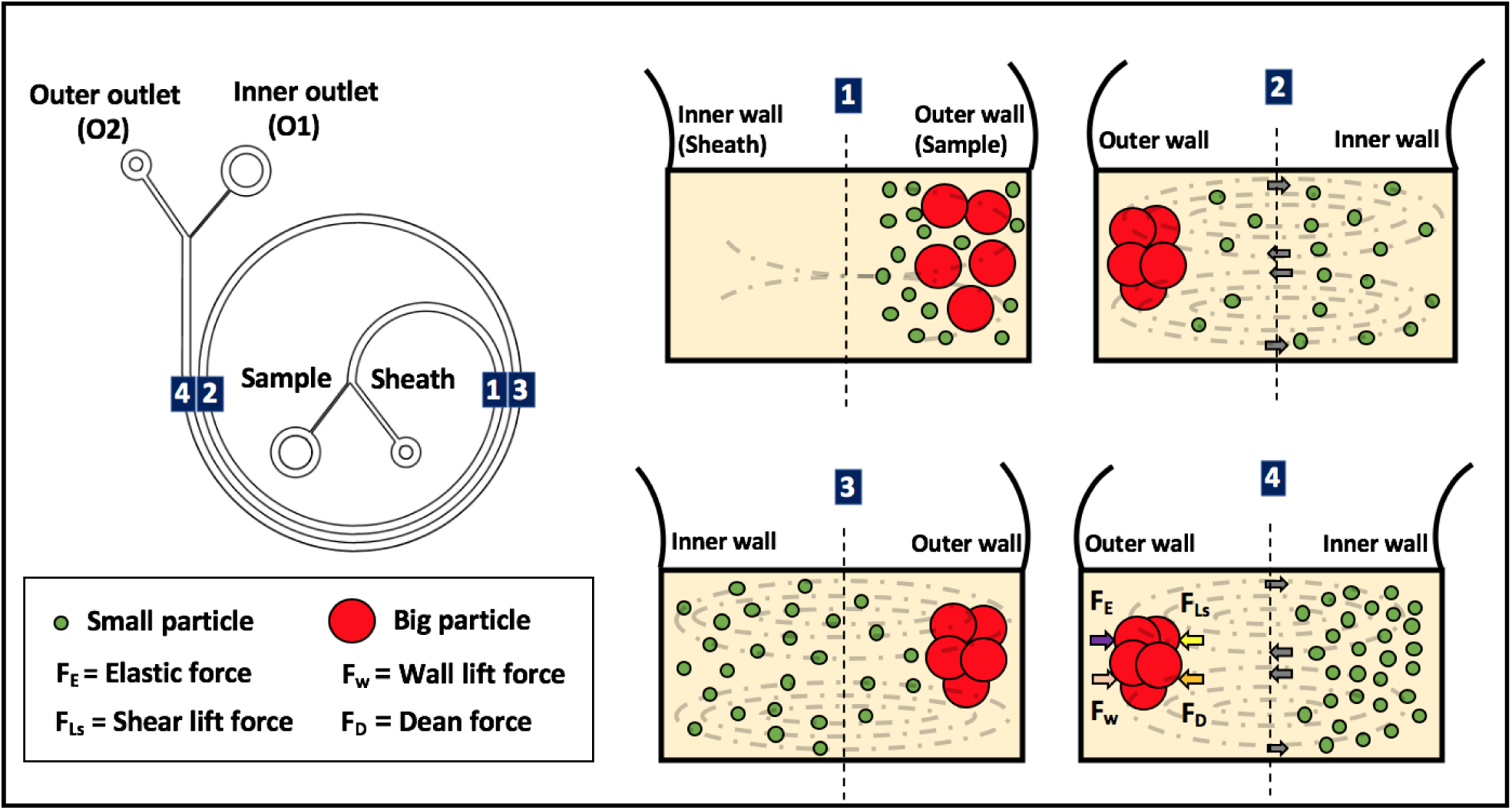
Schematic overview of the bacteria separation method. Particles of two different sizes are shown at different positions (regions 1,2,3 and 4) of a spiral channel. At region 1 (closer to inlet), both particle sizes are pre-positioned at the outer wall by the viscoelastic sheath buffer. As the Dean vortices develop (region 2-4), the smaller particles (green) are dragged with vortices and migrate toward the inner wall while the larger particles (red) remain focused at the outer wall. The larger particles remain focused throughout the channel length due to a balance between the three dominant forces (FL, FE and FD)

### Particle behavior in non-Newtonian fluid

For particle focusing, while scaling differently, all the three forces (F_L_, F_E_ and F_D_) affect the particles and as the forces are fully developed the particles will be fully focused at the equilibrium position close to the outer wall. By carefully optimizing the geometry, flow rate and by prepositioning particles closer to the outer wall at the inlet, we show here that it is possible to achieve focusing and high resolution particle separation. Particles above a size cut-off are fully focused while smaller particles will follow the Dean vortices and differentially migrate towards the inner wall in spiral channels. Fig.2 shows how pre-positioning the particles at the outer wall of the spiral in a viscoelastic flow enables size based particle separation at extremely high volumetric flow rates (1 ml/min). Different sized, 1µm (green) and 7µm (red), particles were suspended in Newtonian (1X PBS) and non-Newtonian (PEO) fluid and prepositioning them at inner and outer wall of the inlet was compared by taking images near the outlet. For Newtonian fluid, it was not possible to separate the particles at a flow rate of 1 mL/min independent of the sample introduction position. In contrast, in a non-Newtonian fluid, starting the sample from the inner wall results in the spreading of 1µm particles toward the center of the channel from the inner wall, while 7 µm migrate to the center of the channel and are unfocused. However, when the sample is introduced from the outer wall, 1 µm particles migrated towards the inner wall while the 7 µm stayed well-focused at the equilibrium position at the outer wall. Clearly, introducing the sample closer to the particle focusing position allows particles above a certain cut-off to be fully focused throughout the entire channel length.

**Fig. 2.**
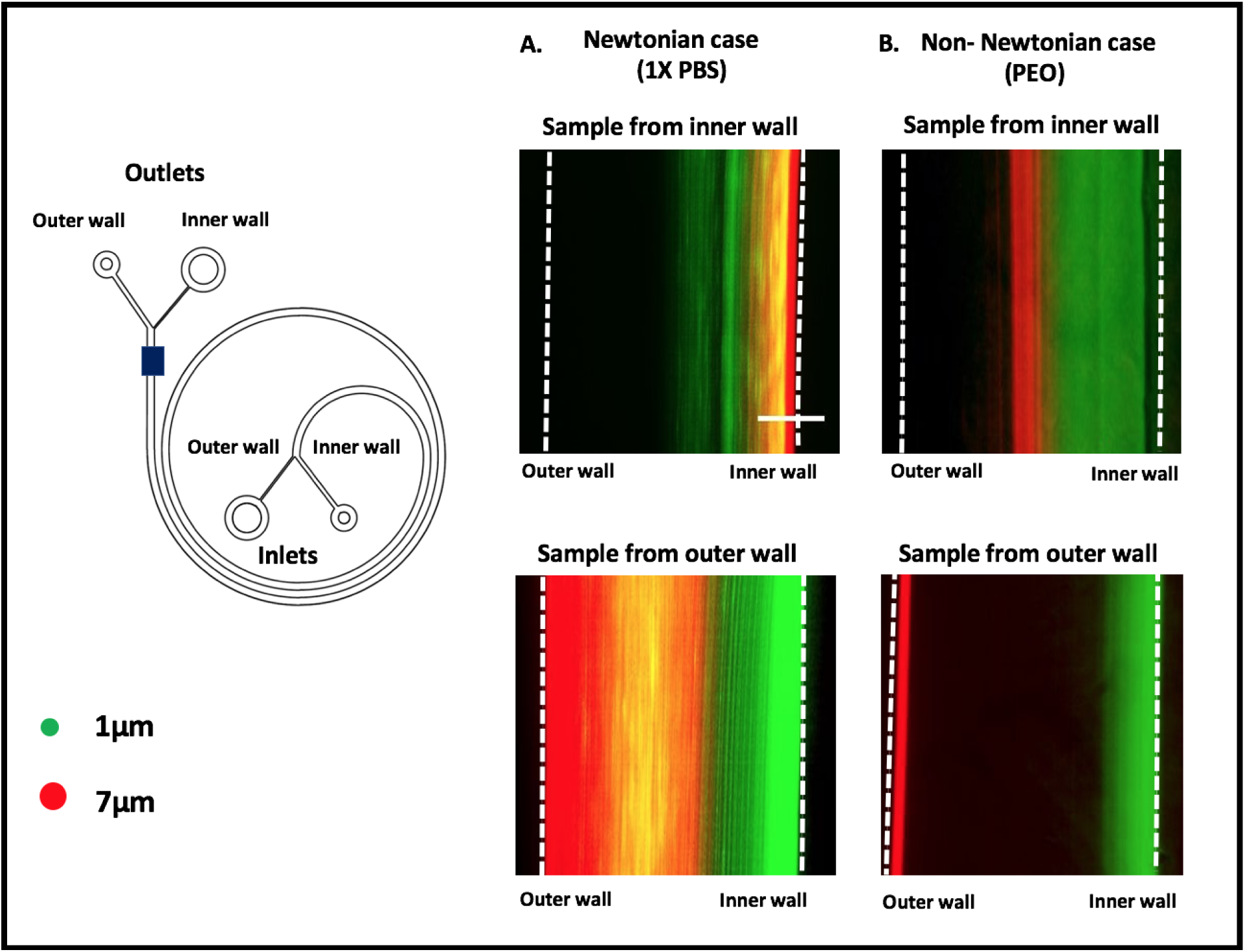
Particle focusing in Newtonian and non-Newtonian fluid. Samples introduced from inner and outer wall of the spiral inlet were studied for (A) Newtonian fluid and (B) Non-Newtonian fluid at a total flow rate of 1 ml/min. Images were taken at the outlets of the spiral (highlighted in blue on the spiral design). The best separation of 1 and 7 µm particles could be obtained using a non-Newtonian fluid when the sample was introduced at the outer wall of the inlet. Scale bar:100 µm.

Following, to better understand particle migration and focusing in a non-Newtonian fluid, a detailed analysis was performed (Fig.3). The sample was introduced at the outer wall, and images were taken at four regions of the spiral channel (region 1-4) in addition to the outlet (region 5) (Fig.3A). Initially, the sheath flow pushes all the particles toward the outer wall (region 1). At region 2, the smaller particles start to migrate towards the inner wall. This is seen in the image as they spread everywhere in the channel. At region 3, most of the smaller particles have reached the inner wall. At region 4, all the smaller particles have now reached the inner wall and stay focused. The smaller particles can then be effectively collected through the inner outlet (region 5). To investigate the influence of Re, the 1 µm and 7 µm particles were pushed at different flow rates and the particle distribution was imaged at the outlet (Fig.3B). The sample flow rate was kept constant at 50 µL/min, while the sheath flow rate was increased systematically from 100 µL/min to 1 mL/min (Re = 5 to 35). As expected, at a low flow rate (Re = 5), both the 1 and 7 µm particles are spread out partly due to insufficient inertial and Dean forces. At relatively low total flow rate there is also insufficient pinching effect of the PEO buffer (1:2 ratio), which will result in the spreading of the particles. As the flow rate increases, a gradual spreading of 1µm was observed and at a flow rate of 500 µL/min (Re = 18), the particles reach to the inner wall. As the flow rate was increased further to 700 µL/min (Re = 25), it was observed that 1 µm particles start to focus toward the inner wall. The particles gradually move close to the inner wall as the flow rate reaches up to 900 µL/min (Re = 32). In contrast, 7 µm particles stay focused toward the outer wall and the higher the flow rate more tightly focused. The particles, initially having a broader distribution of positioning at the outer wall, at flow rates of 100 µL/min (Re = 5), are well focused as the Re was increased (Re =12 – 32). Note that while the Dean forces quickly move the smaller particles laterally toward the inner wall, the migration toward the outer wall in the second round is significantly slowed due to dominant shear-induced lift force counteracting the influence of Dean and elastic forces pushing the particles toward the outer wall again. This phenomenon eventually enables high resolution particle separation.

**Fig.3.**
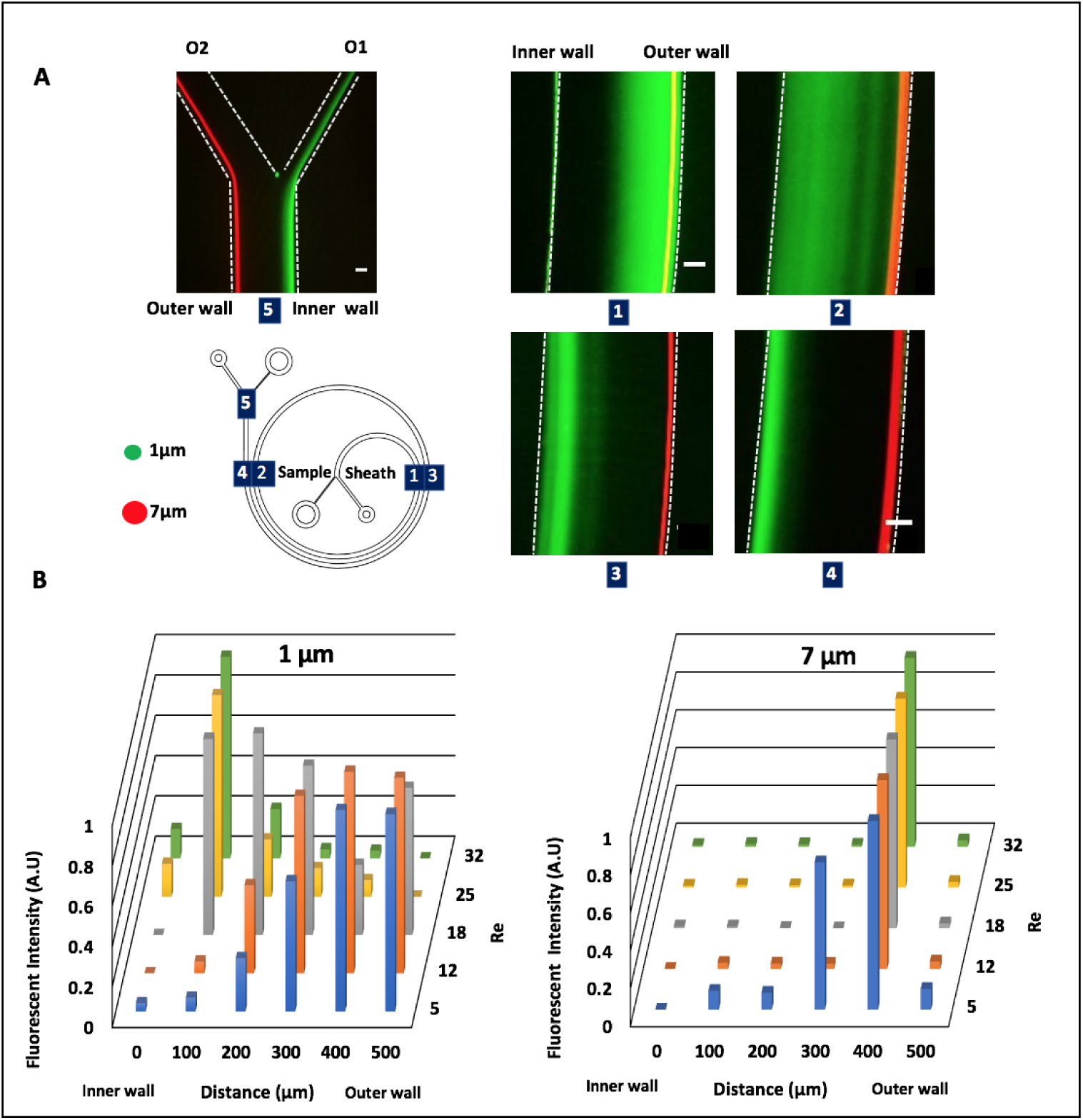
Particle focusing behavior at different Re. 1 µm (green) and 7 µm (in red) particles imaged at five regions of the spiral channel at a total flow rate of 1 mL/min. (A) Near the inlet (region 1), both sized particles are positioned at the outer wall of the channel. The 7 µm particles remain focused at the outer wall in all regions 1-5 while 1 µm particles migrate toward the inner wall (region 4) and can be separated (region 5). Scale bar: 100 µm. (B) Normalized fluorescent cross-section intensity distributions across the channel width at the outlet (region 4). At low Re (Re=5), both 1 and 7µm particles are not focused. At higher flow rates (Re = 25 – 32), the 1 µm particles migrate toward the inner wall, while the 7 µm beads stay focused at the outer wall.

### High resolution particle separation

For sepsis application, the spiral device needs to focus blood cells at the outer wall while allowing the bacteria to migrate away from the wall. To this end, we evaluated the spiral device using particles to find the particle size cut-off above which particles remain focused. Experimentally, 1 µm particles were mixed with particles of different sizes and the migration and focusing was observed (Fig.4A). Using the current spiral design, it is possible to separate 1 µm from 3 µm particles with resolution. While the 2 µm particles are not separated from the 1 µm particles at the outlet, it is noteworthy to mention that while the 1 µm particles have made a turn along the inner wall, the 2 µm particles are yet to make the complete turn. Hence, while outside the scope of this work, by cleverly designing the device it should be possible to differentiate those particles as well. Furthermore, the 3 and µm particles clearly migrate toward the inner wall while the 7 µm particles are fully focused. For a low-aspect-ratio channel geometry (width >> height), focusing is strongly dependent on the particle size to channel height ratio (a/h). Using low-aspect-ratio channel geometry, we previously suggested a minimum a/h ratio > 0.1 for focusing in inertial microfluidics ^49^. Interestingly, our findings also indicate stable focusing at a/h ratio > 0.1. Stable focusing in our case means particles remaining fully focused at the outer wall. This would translate to a particle size above 5 µm in the current spiral design (h=50µm). As a proof of principle for high resolution particle separation, 1 µm and 3 µm particles were mixed and pushed through the device and we analyzed the collected fractions at the outlets (Fig.4B). A hemocytometer was used to count the two fractions. The yield of the 1 μm particles, calculated as fraction of 1 μm particles recovered through the inner outlet (O1) to the total count, was 96%, and the yield was 100% for 3 μm particles in the outer outlet (O2). To reiterate, the total volumetric flow rate used was 1 mL/min, throughput previously only reported for inertial microfluidics.

**Fig.4.**
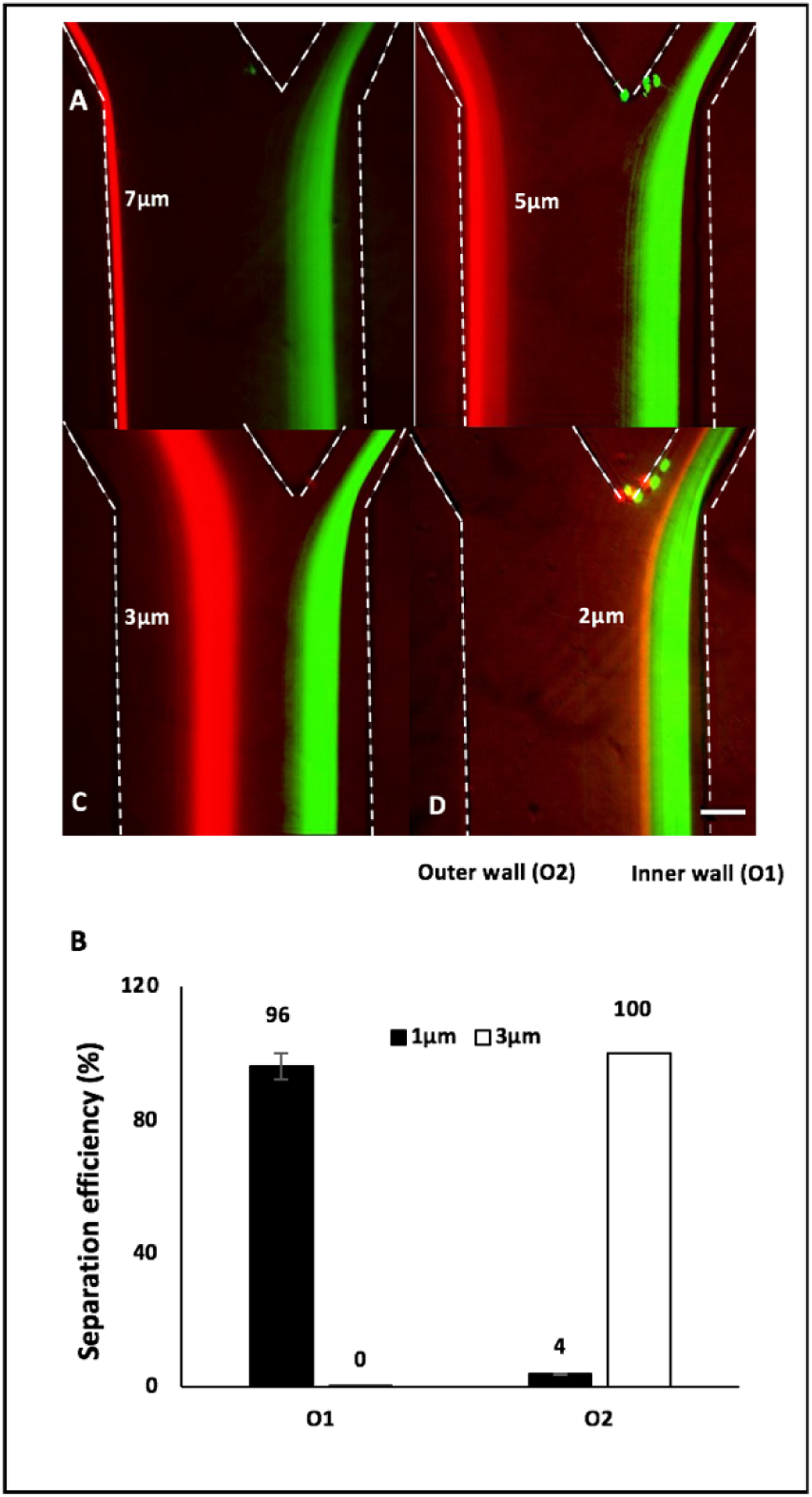
Particle size cut-off. (A) Four different particle sizes (2, 3, 5 and 7 µm shown in red) were mixed with 1 µm particles (in green) in separate experiments and images were taken near the outlet at a flow rate of 1 mL/min. It was observed that as the particle size increases, they get more focused and occupy a stable position toward the outer wall. 3 µm is the smallest particle size that can be readily separated from 1 µm. Scale bar: 100µm. (B) Particle separation efficiency of 1 µm and 3 µm particles. 96% of 1 µm particles were collected at O1 while 100% of 3 µm particles were collected at O2.

### Focusing of blood cells

Blood is a non-Newtonian fluid. The viscosity of the blood is dependent on hematocrit, plasma protein concentration and blood cell count. In flow through spirals, the blood rheology will affect the Dean and lift forces differently. In addition, the visco-elastic nature of the PEO will interact with the blood cells, as well. Initially, we found that whole blood needs to be diluted in order to remain focused. Three different dilutions of blood (1:2, 1:5 and 1:10) were tested. Notably, all the diluted blood cells are focused at the outer wall and effectively collected through the outer outlet channel (Fig.5A). The 1:2 diluted blood (~25% hematocrit) sample is broader, indicating more particle-particle interaction. To investigate the optimal dilution, we mixed 1 µm particles with diluted blood and processed through the spiral device. As shown in Fig.5B, for 1:10 diluted blood (~5% hematocrit), 96% of 1 µm particles were collected at the inner outlet (O1), 93% for 1:5 diluted blood (~10% hematocrit), and 82% for 1:2 diluted blood, respectively. On the other hand, 100% of blood cells were collected in O2 for all three cases. The results indicate the blood cells are fully focused throughout the channel length and as the concentration of blood cells increases, the separation efficiency of 1 µm particles reduces indicating cell-particle interaction. Consequently, there is a need to dilute the blood sample, and an increase in dilution will result in improved separation efficiency of 1 µm particles. It is possible that some particles get stuck to blood cells due to high solid content. In addition, high solid content may ultimately affect how the local forces interact on neighboring particles. Although more work is needed to effectively decipher the effect of solid content, it is evident that high solid content prevents smaller particles from being effectively carried by the Dean vortices toward the inner wall and thus reducing the efficiency of separation. The challenges from particle-particle interaction due to high hematocrit content and the requirement of blood dilution is also reported previously by Shen et al., and Zhou et al., respectively ^50, 51^.

**Fig.5.**
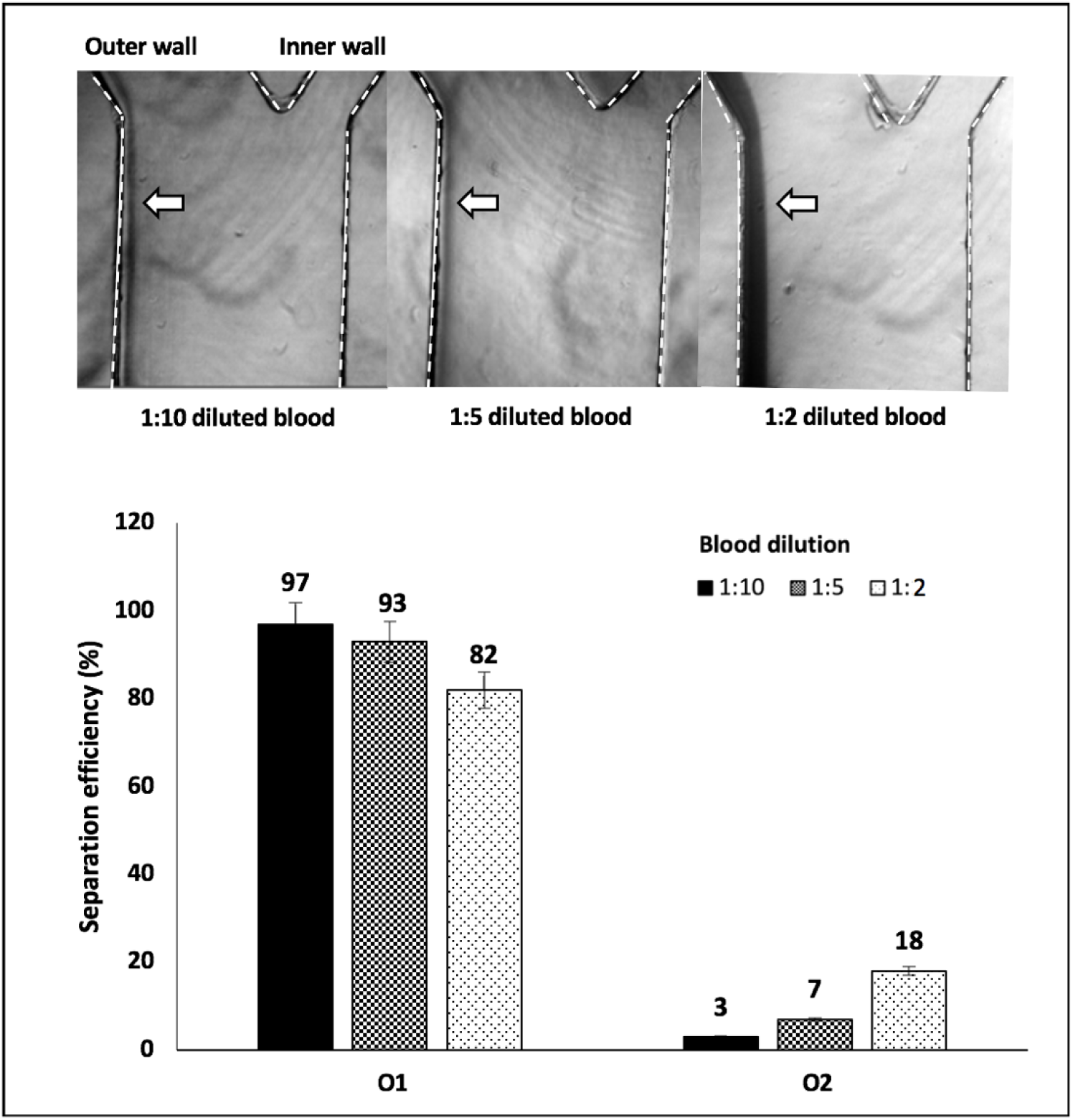
Diluted blood cells focusing. (A) Blood cells remain well focused at the outer wall at higher blood dilutions (1:10 and 1:5). As the hematocrit content increases (25% in case of 1:2 dilution), blood cells begin to spread away from the outer wall due to an increase in particle-particle interaction. Blood dilution is thus important to achieve good focusing and separation of blood cells. Scale bar: 100 µm. (B) Particle separation from diluted blood. Bar graph showing the separation of 1µm particles with an efficiency of 97%, 93%, and 83% from 1:10, 1:5, and 1:2 diluted blood, respectively.

**Fig.5.**
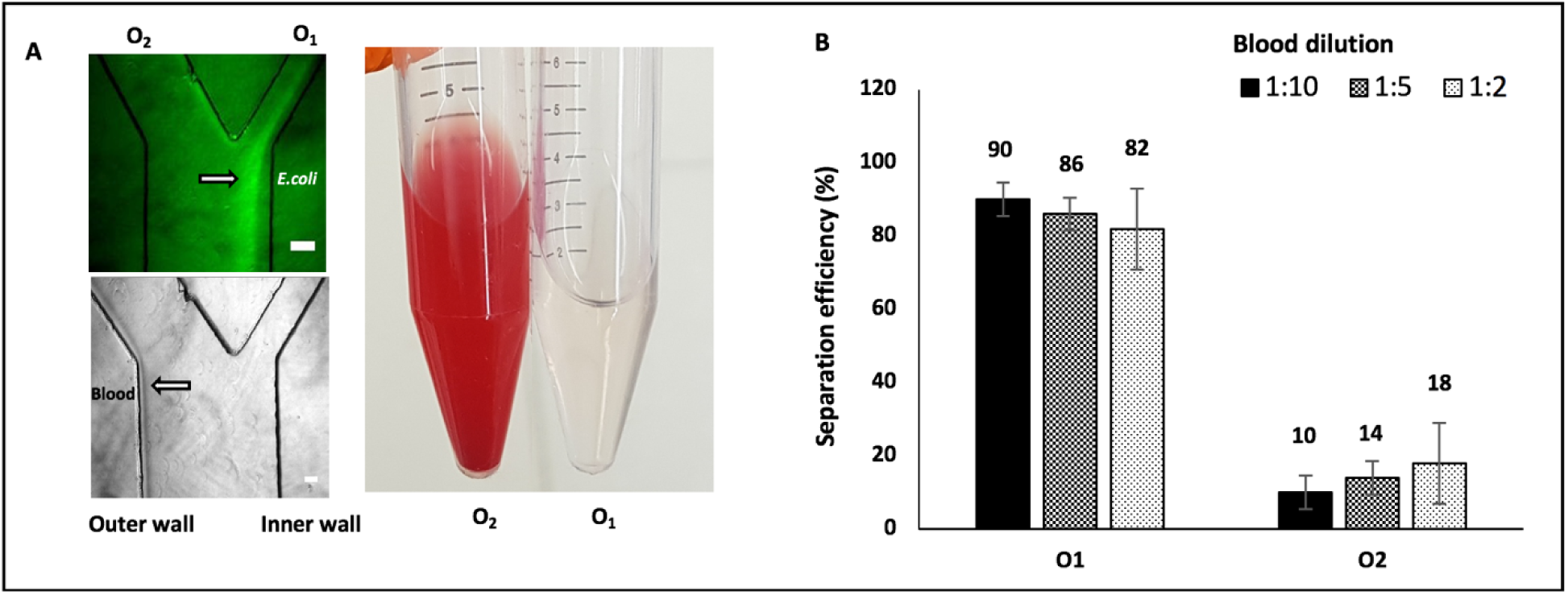
*E.coli* separation from diluted blood. (A) Qualitative analysis showing fluorescently labelled (using bacterial viability kit) *E.coli* migrating to the inner wall and collected in O1 while blood cells remain focused at the outer wall and collected in O2. Scale bar: 100 µm. (B) Quantification of *E.coli* (1000 CFU/mL) spiked into three diluted blood samples (1:10, 1:5, and 1:2). Agar plating based quantification show *E.coli* separation efficiency between 90% to 82% depending on blood dilution.

### Bacteria separation from diluted blood

Before testing the behavior of bacteria spiked in blood, we tested how bacteria behave in viscoelastic fluids. To demonstrate this, PEO was spiked with fluorescently labelled gram negative *Escherichia coli* (*E.coli*) and gram positive *Staphylococcus capitis (Staph)* (10^7^ CFU/mL) processed through the spiral device at a total flow rate of 1mL/min in separate experiments. We observed a similar behavior for both gram negative and gram positive with an yield of 94% and 89% for *E.coli* and *Staph* respectively using agar plating based quantification (supplementary Fig.S1). Following, to study the separation of blood cells, lower concentration of *E.coli* (10^3^ CFU/mL) was spiked into 1:10, 1:5, and 1:2 diluted blood. As expected, the bacteria migrated to the inner wall and were collected at outlet O1, whereas blood cells stay near the outer wall and collected at outlet O2 (Fig.5A). Quantification performed using blood agar plating showed a separation efficiency of 82 to 90% of *E.coli* (Fig.5B) depending on the dilution, while all blood cells were recovered through the outer outlet for all cases. The higher blood dilution, the higher bacteria separation efficiency, in agreement with the results obtained for 1µm particles (see Fig.4). While high throughput is desirable for any application, need to process large volumes is extremely very important for sepsis applications. Ideally, this should not be on the cost of efficiency. Using our current spiral device, it takes 40 minutes to process 1 mL of whole blood at a separation efficiency of 82%. To the best of our knowledge, this is the highest throughput yet reported for a single device at an efficiency above 80%. On the other hand, for a separation efficiency of 90%, it would take approximately three hours to process 1 mL blood.

For sepsis applications, ideally large volumes (2-7 mL whole blood) will be necessary for any sample preparation method. Microfluidics, defined as the science and technology of systems that can handle small volumes of fluid in channels of ten to few hundred microns in size, has proven to be well suited for applications in clinical diagnostics. However, while impressive progress has been made in high throughput rare circulating tumor cell isolation, there has been limited success for bacteria isolation from blood using microfluidics. In general, there is a trade of between separation efficiency and throughput for any given technology. To this end, in comparison to the current state of the art, our method is well suited for high separation efficiency without compromising throughput. In Fig.6, we compare work published using both active (magnetophoresis ^38^, acoustophoresis ^32, 33^ and dielectrophoresis ^35, 36^) and passive (size-based separation ^28, 31, 34, 37^) to our current work for bacteria isolation from blood. Most of the work in inertial microfluidics that show good separation efficiency have lower throughput ^28–36^ while studies that highlight higher throughput show lower separation efficiency ^37, 38^. The separation efficiency corresponding to the three different blood dilutions in the current work is highlighted in red in Fig.6. Using a single spiral, we demonstrate processing of 1 mL of blood at extremely high throughput with separation efficiency >80% for all the blood dilutions tested. To our best knowledge, this is the highest throughput reported for a single passage using a single chip at the given high separation efficiency.

**Fig.6.**
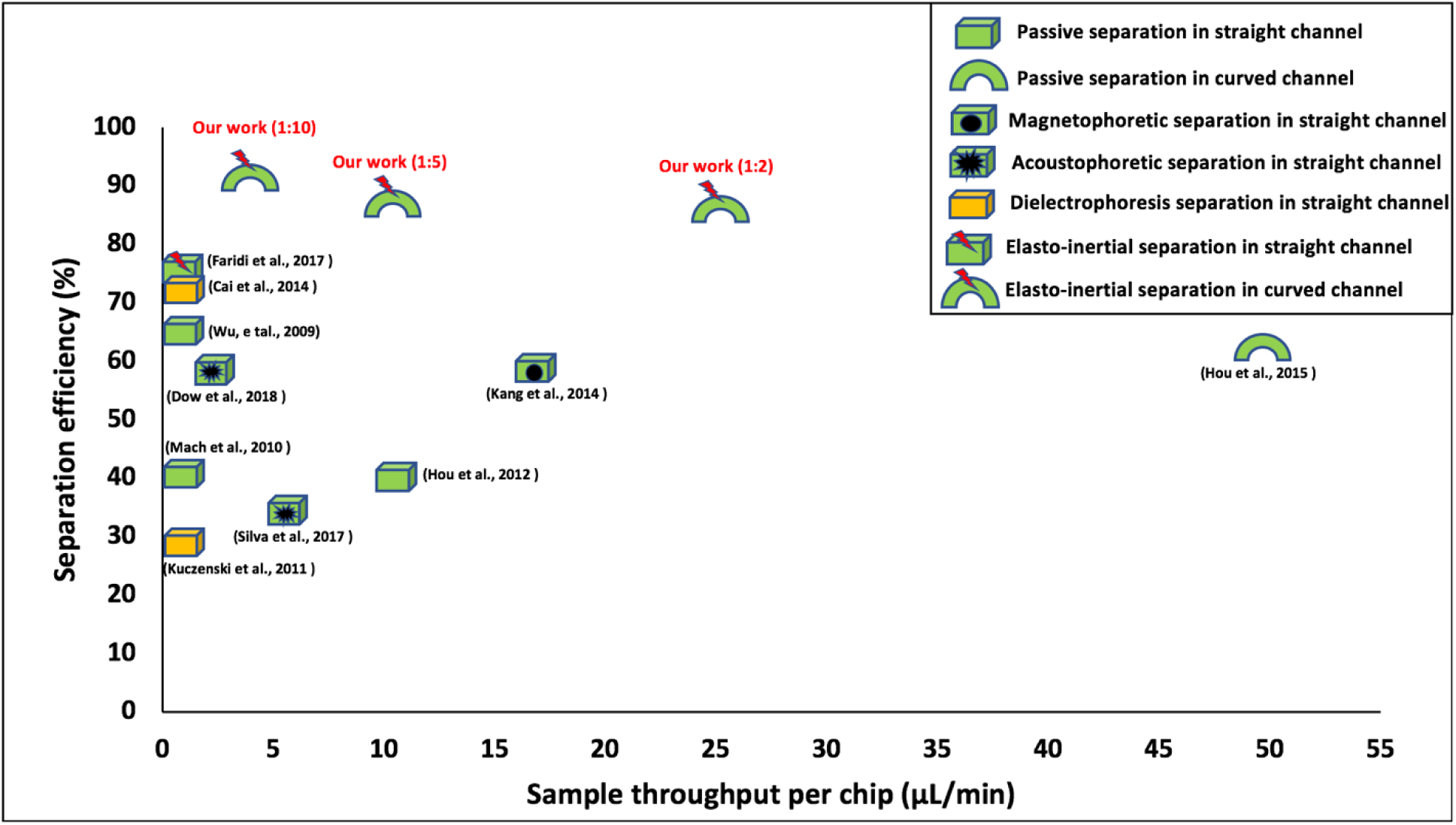
Summary of current state of art microfluidics-based bacteria separation methods from blood. In general, there is a trade-off between separation efficiency and sample throughput using both active and passive methods. Our current work for the different diluted blood samples is shown in red having the highest efficiency (> 80%) using single passage, single chip and passive method based elasto-inertial microfluidics.

One way to increase the throughput in our current setup is via parallelization. We have previously presented on a highly scalable, lithography-defined microfabrication method for passive size-based particle separation^52^. For spiral design, it is straightforward to stack the devices vertically, as reported by Warkiani., et al ^53^. As a proof of principle, we tested stacking two spiral channels and tested particle separation (supplementary Fig.S3). At a total flow rate of 3 mL/min, complete separation of 1µm and 7µm particles is shown in supplementary Fig.S3. Using only four spirals stacked, and without any further optimization, a throughput of 6 mL of whole blood can be processed within an hour for 1:2 diluted blood at a separation efficiency of 90%. Future studies should focus on expanding the bacteria strains, including common pathogenic strains for clinical implementation. While outside the scope of this paper, we have initiated the work on stacked spiral devices for clinical implementation of the stand-alone sample preparation module. We anticipate the method presented here to be of high value in clinical sample preparation applications for both nucleic acid based molecular diagnostics and phenotypic analysis.

## Conclusion

In this paper, we present a sample preparation module using elasto-inertial microfluidics based on passive separation method, consisting of a two turn spiral channel design for separating viable bacteria from blood at high throughput and efficiency. In this work, we use viscoelastic buffer to sheath the blood sample to the outer wall of a spiral and demonstrate for the first time that blood cells remain fully focused at the outer wall throughout the entire spiral length while bacteria continuously migrate due to the dean vortices for efficient separation. The viable bacteria are recovered free of blood cells and readily available for downstream analysis. This not only show that the larger particles are fully focused at the outer wall but the importance of F_E_ enabling high resolution focusing and separation. In addition, we address one of the main challenges of miniaturization for sepsis application: the trade-off between separation efficiency and throughput. Here, we demonstrate separation of *E.coli* from 1mL of blood in 40 minutes at an efficiency of 82%. The microfluidic platform opens up possibilities of effectively and rapidly separating viable bacteria from blood at throughputs unmatched using existing microfluidics methods and should warrant clinical value as a stand-alone sample preparation method in clinical settings.

## Materials and methods

### Fabrication of the microchannels

The master mold was obtained using standard photolithography techniques. AutoCAD 2017, (AutoDesk, USA) software was used to design the two-turn spiral channel. It was printed on a Mylar mask. A negative photoresist SU-8 was spin-coated on to a 4-inch silicon wafer. It was exposed to a UV light source through the Mylar mask and was developed using SU-8 developer to obtain the channel design onto the silicon wafer that was ordered from micro resist technology, Germany. A PDMS replica was produced using a soft lithography technique where Sylgard 184 silicon elastomer and curing agent was mixed in a 10:1 ratio. The silicon elastomer was poured onto the SU-8 master and was cured for a minimum. of 6 hr. at 65°C. The PDMS mold was cut and holes were punched using a 0.75 mm diameter Harris Uni-Core puncher to obtain inlets and outlets. Clean glass and the PDMS mold was bonded using an oxygen plasma treatment using FEMTO SCIENCE CUTE plasma system, South Korea and incubation at 110°C for 10 min to ensure robust bonding.

### Chip characterization

A two-turn spiral channel with a rectangular cross-sectional geometry was first designed and tested by Bhagat et al. ^54^ to isolate CTCs from blood. A similar design was used to study the behavior of particles of different sizes in our work. The design is shown in Fig. 1 in the center. The microchannel had a width of *w* = 500 µm and a height of *h* = 50 µm. The aspect ratio, defined as width/height (w/h) was 10. The blockage ratio, which is defined as a/h, where ‘a’ is the diameter of the particle and ‘h’ is the channel height, ranged from 0.02 to 0.14.

### Particle experiments

Different diameters of fluorescently labelled, neutrally buoyant polystyrene particles from ThermoFischer Scientific of 10µL were spiked into 10 mL of 150 ppm concentration polyethylene oxide (PEO), Molecular weight = 2,000 kDa and was used as a sample fluid. PEO of same concentration without polystyrene particles were used as sheath fluid. Different sized particles such as 1, 3, 5 and 7µm from the same company were used for particle experiments. A Harvard Apparatus Pump 33 DDS syringe pump was used to pump the sample and sheath flowing from a 20 mL BD plastic syringes. The syringes were connected to the inlets and outlets of the chip using COLE-PARMER TYGON tubing with inner diameters of 0.254 mm and outer diameter 0.762 mm. The migration and focusing of particles in the microchannels was observed using AXIOVERT 135 TV inverted fluorescence microscope. The sample flow rate for all experiments was kept at 50 µL/min. The sheath flow rate was varied based on the experiment. Fluorescent quantification was performed using ImageJ freeware (https://imagej.nih.gov/ij/).

### Blood and bacteria-based experiments

Healthy blood samples were collected from the blood center GeBlod, Stockholm, Sweden in EDTA tubes. PEO was used to dilute blood in 1:10, 1:5, and 1:2 dilutions. Inlets and outlets were centrifuged at 5000 g for 10 min to pellet bacteria and resuspended into equal volumes of PEO. *Escherichia coli* ATCC 25922 strain (*E.coli*) and *Staphylococcus capitis (Staph)* was grown separately on blood agar plates (3 sets of plates from each outlet per experiment) for 12 hr. at 37 °C incubator. Optical density was measured using an Ultrospec 10 cell density meter (Amersham Biosciences, UK) at 600 nm after inoculating bacteria from the blood agar plate into PEO. Bacteria were fluorescently labelled using the Backlight viability kit (ThermoFischer, Sweden) before spiking into blood for the purpose of visualization and to check their viability. Green fluorescence depicts viable bacteria while red fluorescence indicates dead ones.

## Acknowledgment

This project has received funding from the European Union Horizon 2020 research and innovation programme under the Marie Skłodowska-Curie grant agreement No 675412 as a part of the consortium New diagnostics for infectious diseases (ND4ID).

## References

1. H. Li, L. Liu, D. Y. Zhang, J. Y. Xu, H. P. Dai, N. Tang, X. Su and B. Cao, Lancet, 2020, 395, 1517–1520.

2. M. Singer, C. S. Deutschman, C. W. Seymour, M. Shankar-Hari, D. Annane, M. Bauer, R. Bellomo, G. R. Bernard, J. D. Chiche, C. M. Coopersmith, R. S. Hotchkiss, M. M. Levy, J. C. Marshall, G. S. Martin, S. M. Opal, G. D. Rubenfeld, T. van der Poll, J. L. Vincent and D. C. Angus, JAMA, 2016, 315, 801–810.

3. F. B. Mayr, S. Yende and D. C. Angus, Virulence, 2014, 5, 4–11.

4. J. Cohen, P. Cristofaro, J. Carlet and S. Opal, Critical Care Medicine, 2004, 32, 1510–1526.

5. J. L. Vincent, J. Rello, J. Marshall, E. Silva, A. Anzueto, C. D. Martin, R. Moreno, J. Lipman, C. Gomersall, Y. Sakr, K. Reinhart and E. I. G. Investigators, Jama-J Am Med Assoc, 2009, 302, 2323–2329.

6. C. Fleischmann, A. Scherag, N. K. J. Adhikari, C. S. Hartog, T. Tsaganos, P. Schlattmann, D. C. Angus, K. Reinhart and I. F. A. C. Trialists, Am J Resp Crit Care, 2016, 193, 259–272.

7. WHO, Improving the prevention, diagnosis and clinical management of sepsis, https://www.who.int/servicedeliverysafety/areas/sepsis/en/).

8. Journal, 2020.

9. K. E. Rudd, S. C. Johnson, K. M. Agesa, K. A. Shackelford, D. Tsoi, D. R. Kievlan, D. V. Colombara, K. S. Ikuta, N. Kissoon, S. Finfer, C. Fleischmann-Struzek, F. R. Machado, K. K. Reinhart, K. Rowan, C. W. Seymour, R. S. Watson, T. E. West, F. Marinho, S. I. Hay, R. Lozano, A. D. Lopez, D. C. Angus, C. J. L. Murray and M. Naghavi, Lancet, 2020, 395, 200–211.

10. C. J. Paoli, M. A. Reynolds, M. Sinha, M. Gitlin and E. Crouser, Crit Care Med, 2018, 46, 1889–1897.

11. C. E. Edmiston, R. Garcia, M. Barnden, B. DeBaun and H. B. Johnson, Am J Infect Control, 2018, 46, 1060–1068.

12. A. Kumar, D. Roberts, K. E. Wood, B. Light, J. E. Parrillo, S. Sharma, R. Suppes, D. Feinstein, S. Zanotti, L. Taiberg, D. Gurka, A. Kumar and M. Cheang, Crit Care Med, 2006, 34, 1589–1596.

13. U. Frank, D. Malkotsis, D. Mlangeni and F. D. Daschner, Eur J Clin Microbiol, 1999, 18, 248–255.

14. J. A. Garcia-Prats, T. R. Cooper, V. F. Schneider, C. E. Stager and T. N. Hansen, Pediatrics, 2000, 105, 523–527.

15. Y. Haimi-Cohen, E. M. Vellozzi and L. G. Rubin, J Clin Microbiol, 2002, 40, 898–901.

16. A. Mobed, B. Baradaran, M. de la Guardia, M. Agazadeh, M. Hasanzadeh, M. A. Rezaee, J. Mosafer, A. Mokhtarzadeh and M. R. Hamblin, Trac-Trend Anal Chem, 2019, 113, 157–171.

17. M. Sinha, J. Jupe, H. Mack, T. P. Coleman, S. M. Lawrence and S. I. Fraley, Clin Microbiol Rev, 2018, 31.

18. K. S. Gracias and J. L. McKillip, Can J Microbiol, 2004, 50, 883–890.

19. C. W. Yung, J. Fiering, A. J. Mueller and D. E. Ingber, Lab Chip, 2009, 9, 1171–1177.

20. S. Q. Wang, F. Inci, T. L. Chaunzwa, A. Ramanujam, A. Vasudevan, S. Subramanian, A. C. F. Ip, B. Sridharan, U. A. Gurkan and U. Demirci, Int J Nanomed, 2012, 7, 2591–2600.

21. J. J. Lee, K. J. Jeong, M. Hashimoto, A. H. Kwon, A. Rwei, S. A. Shankarappa, J. H. Tsui and D. S. Kohane, Nano Lett, 2014, 14, 1–5.

22. D. Di Carlo, D. Irimia, R. G. Tompkins and M. Toner, P Natl Acad Sci USA, 2007, 104, 18892–18897.

23. X. Y. Ding, Z. L. Peng, S. C. S. Lin, M. Geri, S. X. Li, P. Li, Y. C. Chen, M. Dao, S. Suresh and T. J. Huang, P Natl Acad Sci USA, 2014, 111, 12992–12997.

24. X. Y. Hu, P. H. Bessette, J. R. Qian, C. D. Meinhart, P. S. Daugherty and H. T. Soh, P Natl Acad Sci USA, 2005, 102, 15757–15761.

25. A. Ashkin, Physical Review Letters, 1970, 24, 156–&.

26. N. Pamme, Lab Chip, 2006, 6, 24–38.

27. C. Liu and G. Q. Hu, Micromachines-Basel, 2017, 8.

28. Z. G. Wu, B. Willing, J. Bjerketorp, J. K. Jansson and K. Hjort, Lab Chip, 2009, 9, 1193–1199.

29. S. H. Jung, Y. K. Hahn, S. Oh, S. Kwon, E. Um, S. Choi and J. H. Kang, Small, 2018, 14.

30. M. A. Faridi, H. Ramachandraiah, I. Banerjee, S. Ardabili, S. Zelenin and A. Russom, J Nanobiotechnology, 2017, 15, 3.

31. A. J. Mach and D. Di Carlo, Biotechnol Bioeng, 2010, 107, 302–311.

32. R. Silva, P. Dow, R. Dubay, C. Lissandrello, J. Holder, D. Densmore and J. Fiering, Biomed Microdevices, 2017, 19.

33. P. Dow, K. Kotz, S. Gruszka, J. Holder and J. Fiering, Lab Chip, 2018, 18, 923–932.

34. H. W. Hou, H. Y. Gan, A. A. S. Bhagat, L. D. Li, C. T. Lim and J. Han, Biomicrofluidics, 2012, 6.

35. D. Y. Cai, M. Xiao, P. Xu, Y. C. Xu and W. B. Du, Lab Chip, 2014, 14, 3917–3924.

36. R. S. Kuczenski, H. C. Chang and A. Revzin, Biomicrofluidics, 2011, 5.

37. H. W. Hou, R. P. Bhattacharyya, D. T. Hung and J. Han, Lab Chip, 2015, 15, 2297–2307.

38. J. H. Kang, M. Super, C. W. Yung, R. M. Cooper, K. Domansky, A. R. Graveline, T. Mammoto, J. B. Berthet, H. Tobin, M. J. Cartwright, A. L. Watters, M. Rottman, A. Waterhouse, A. Mammoto, N. Gamini, M. J. Rodas, A. Kole, A. Jiang, T. M. Valentin, A. Diaz, K. Takahashi and D. E. Ingber, Nat Med, 2014, 20, 1211–1216.

39. M. E. Warkiani, B. L. Khoo, L. D. Wu, A. K. P. Tay, A. A. S. Bhagat, J. Han and C. T. Lim, Nature Protocols, 2016, 11, 134–148.

40. N. Xiang, X. J. Zhang, Q. Dai, J. Cheng, K. Chen and Z. H. Ni, Lab Chip, 2016, 16, 2626–2635.

41. S. Yang, J. Y. Kim, S. J. Lee, S. S. Lee and J. M. Kim, Lab Chip, 2011, 11, 266–273.

42. G. D’Avino, G. Romeo, M. M. Villone, F. Greco, P. A. Netti and P. L. Maffettone, Lab Chip, 2012, 12, 1638–1645.

43. G. Segre and A. Silberberg, Nature, 1961, 189, 209–&.

44. A. A. S. Bhagat, S. S. Kuntaegowdanahalli and I. Papautsky, Phys Fluids, 2008, 20.

45. S. A. Berger, L. Talbot and L. S. Yao, Annu Rev Fluid Mech, 1983, 15, 461–512.

46. A. M. Leshansky, A. Bransky, N. Korin and U. Dinnar, Phys Rev Lett, 2007, 98, 234501.

47. B. P. Ho and L. G. Leal, J Fluid Mech, 1976, 76, 783–799.

48. H. Giesekus, Rheol Acta, 1982, 21, 366–375.

49. A. Russom, A. K. Gupta, S. Nagrath, D. Di Carlo, J. F. Edd and M. Toner, New J Phys, 2009, 11, 75025.

50. S. F. Shen, C. Ma, L. Zhao, Y. L. Wang, J. C. Wang, J. Xu, T. B. Li, L. Pang and J. Y. Wang, Lab Chip, 2014, 14, 2525–2538.

51. J. Zhou, P. V. Giridhar, S. Kasper and I. Papautsky, Lab Chip, 2013, 13, 1919–1929.

52. J. Hansson, J. M. Karlsson, T. Haraldsson, H. Brismar, W. van der Wijngaart and A. Russom, Lab Chip, 2012, 12, 4644–4650.

53. M. E. Warkiani, A. K. P. Tay, G. F. Guan and J. Han, Sci Rep-Uk, 2015, 5.

54. A. A. S. Bhagat, S. S. Kuntaegowdanahalli and I. Papautsky, Lab Chip, 2008, 8, 1906–1914.

