## Supplementary data for "High resolution and high throughput bacteria separation from blood using elasto-inertial microfluidics"

### Table of Contents

1. **Bacteria focusing in PEO buffer**
2. **Quantification of outlets by blood agar plating**
3. **Increased throughput by parallelization approach**

**Bacteria focusing in PEO buffer**

*E.coli* (gram negative) and *Staphylococcus* (gram positive) (both at concentration of $10^7$ CFU/mL) were spiked into PEO buffer in two separate experiments and separated fractions were analyzed. This is shown in Fig.S1. Quantification using blood agar plating (n=3) showed that an yield of 94% of *E.coli* and 89% of *Staphylococcus* was obtained at O1.

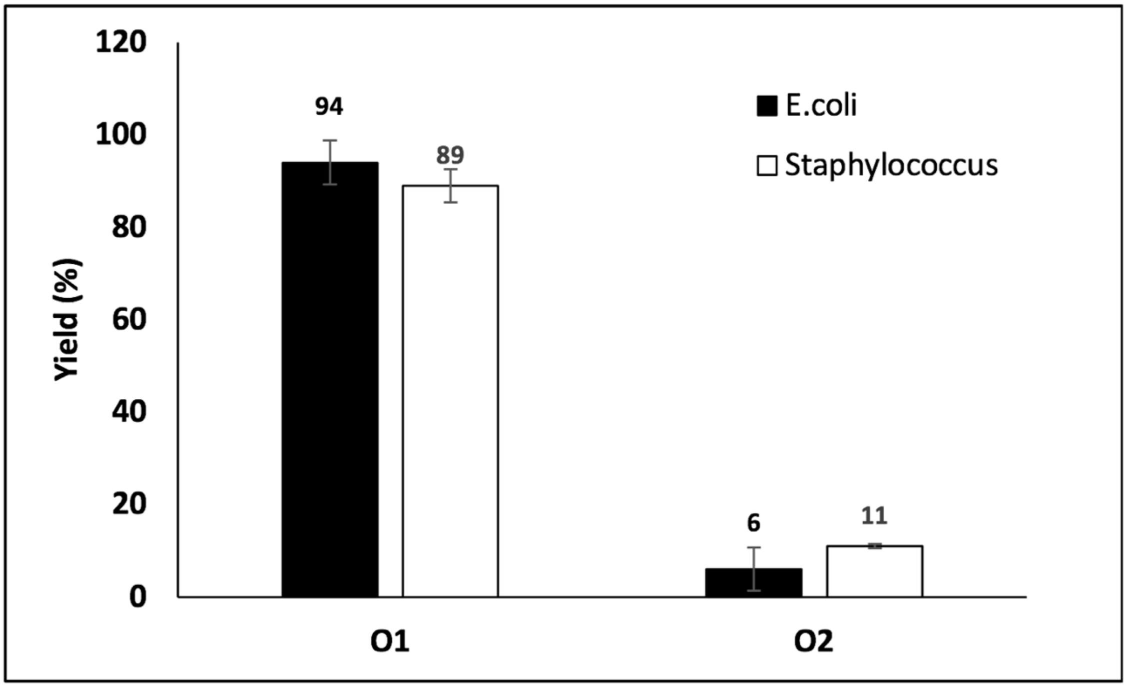


**Fig.S1**. **Bacteria focusing**. *E.coli* and *Staphylococcus* ($10^7$ CFU/mL) spiked into PEO buffer in separate experiments and sample was processed through the spiral device at a total flow rate of 1 mL/min and the collected fractions were analyzed. Bacteria were collected at the desire (inner wall, O1) outlet at a yield of 94% for *E.coli* and 89% for *Staphylococcus* respectively (n=3).

**Quantification of outlets by blood agar plating**

Both inlet and outlets (O1 and O2) after the separation of *E.coli* from different blood dilutions (1:10, 1:5 and 1:1) were plated on a blood agar plate to compare and quantify *E.coli* growth. Three experiments for each dilution were performed and three sets of blood agar plates for each case were plated, but only one set for each is shown here in Fig. S2. The bacterial colonies were counted from the plates for quantification.

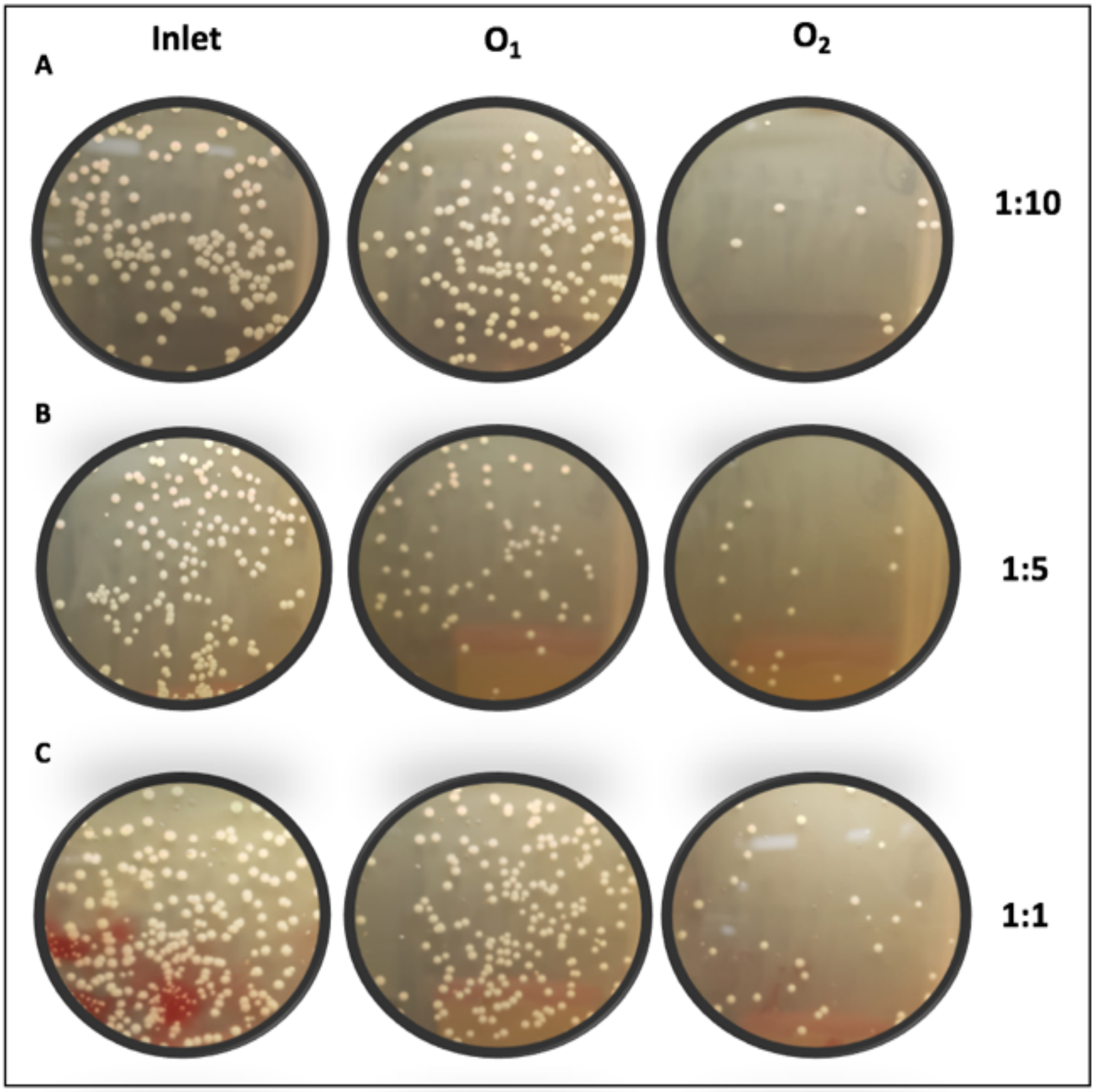


**Fig.S2**. **Quantification of E.coli separated from blood using blood agar plating**. Inlet and outlets (O1 and O2) plated on blood agar after *E.coli* separation from different blood dilutions 1:10, 1:5, and 1:1 to compare and quantify *E.coli* growth. Quantification was performed by counting the bacterial colonies.

**Parallelization for increased throughput**

To demonstrate the possibility of processing high volumes of blood, using parallelization two spiral chips were stacked vertically (Fig.S3). Using the same experimental conditions as before, it was observed that 1µm (green) and 7µm (red) could be separated at a very high throughput of 3mL/min.

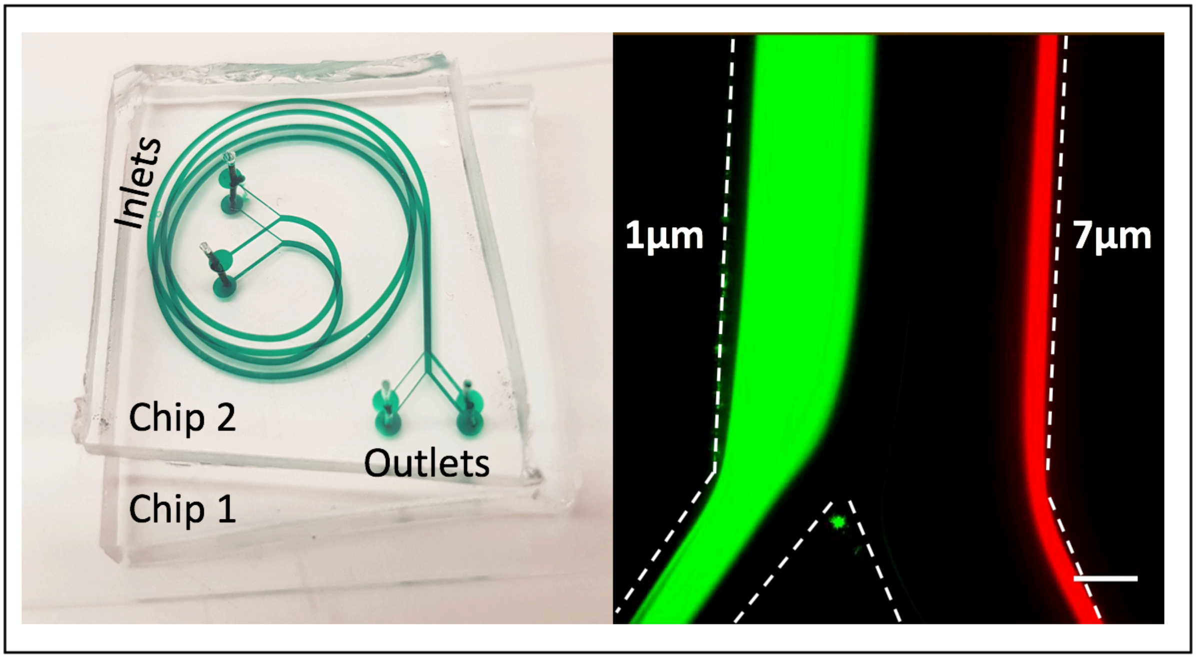


**Fig.S3**. Parallelization approach achieves higher throughput. (Left) Two spiral chips were vertically aligned and bonded to produce a two-layer parallel spiral device (Right). Chip characterization. 1µm and 7µm particles were separated at a total flow rate of 3 mL/min showing the potential of achieving higher throughput by parallelization.
